# Transcriptome-wide analysis of alternative splicing in women with fibromyalgia highlights GNLY as an immune candidate

**DOI:** 10.64898/2026.09.08.750016

**Authors:** Yannick Bruns, Alexander Neumann

**Affiliations:** Department of Anaesthesiology and Intensive Care Medicine, Pain Clinic, Hannover Medical School, Hannover, Germany; Institute for Neuropathology, University Medical Center Göttingen, Göttingen, Germany

**Author notes:** **Correspondence:** Dr. med. Yannick Bruns, Klinik für Anästhesiologie und Intensivmedizin, Schmerzambulanz Medizinische Hochschule Hannover, Carl-Neuberg-Straße 1, 30625 Hannover, Germany.

**Keywords:** fibromyalgia, alternative splicing, LeafCutter, RNA-seq, peripheral blood, granulysin, cell composition deconvolution

## Abstract

**Background:** Fibromyalgia syndrome (FMS) affects 1 to 2 percent of the general population, with higher prevalence estimates in women. Recent evidence supports peripheral immune involvement in disease pathophysiology. Alternative splicing (AS) regulates immune cell function independently of transcript abundance and carries disease-relevant signal in related autoimmune conditions. No study has performed transcriptome-wide differential AS analysis directly in RNA-sequencing data from individuals with FMS.

**Methods:** We performed a secondary analysis of GSE221921, restricted to female participants (n = 91 FMS, n = 41 controls). Peripheral immune cell composition was estimated with ABIS, a deconvolution method specifically trained on peripheral blood mononuclear cell signatures. Differential AS was analysed with LeafCutter incorporating cell composition principal components as covariates. Differential gene expression was performed with DESeq2 under the same adjustment framework.

**Results:** FMS samples showed lower ABIS-inferred conventional monocyte estimates (Cliff’s δ = - 0.38, FDR = 0.008) and higher memory B-cell estimates (Cliff’s δ = +0.32, FDR = 0.030). Composition-adjusted LeafCutter analysis identified 14 significant intron clusters, of which 9 were retained for biological interpretation. GNLY (granulysin) was the primary candidate, with a composition-adjusted 5’ junction usage shift illustrated by pooled Sashimi visualisation. Composition adjustment substantially reduced significant differentially expressed genes from 10,803 to 3,186 (baseMean ≥ 10).

**Conclusions:** This first transcriptome-wide differential AS analysis in FMS identifies GNLY, encoding the cytolytic lymphocyte effector granulysin, as a splicing-specific candidate for functional follow-up. AS captures FMS-associated signal not detectable by gene-level expression analysis, and composition adjustment substantially altered the differential expression landscape.

## 1. Introduction

Fibromyalgia syndrome (FMS) affects approximately 1 to 2 percent of the general population, with higher prevalence estimates in women (D’Souza et al., 2026; Heidari et al., 2017). It remains one of the most disabling chronic pain conditions (Häuser et al., 2015). Patients present with widespread pain, fatigue, sleep disturbance, and cognitive dysfunction, and current pharmacological options provide limited relief in most cases (Häuser et al., 2018). The molecular basis of the disease is poorly understood, and diagnosis still relies entirely on clinical criteria (Wolfe et al., 2016).

Recent large-scale genetic studies have established the polygenic architecture of FMS and predominantly implicate neuronal biology (Kerrebijn et al., 2026; Bright et al., 2026). In parallel, experimental and immunological work has provided evidence for peripheral immune involvement, including passive transfer of pain-related phenotypes by patient IgG and disrupted peripheral B cell tolerance (Goebel et al., 2021; Long et al., 2025). Peripheral blood transcriptomics addresses peripheral immunity directly, but comprehensive analysis requires attention to alternative splicing (AS) as well as to expression, and to the cell-composition heterogeneity that shapes both.

Two published transcriptomic studies have analysed the largest available FMS RNA-sequencing dataset, GSE221921 (96 FMS patients, 93 controls, PBMCs). The original study identified 1,720 differentially expressed genes and defined three transcriptomic subgroups (FM1, FM2, FM3) with distinct enrichment patterns (Mohapatra et al., 2024). A subsequent reanalysis derived an XGBoost machine-learning classifier from FMS-associated differential expression across multiple cohorts, identifying three candidate diagnostic biomarkers (DYRK3, RGS17, ARHGEF37); the same study estimated immune-cell proportions using CIBERSORT (Zhao et al., 2025). Neither study examined AS, and neither incorporated inferred cell composition as covariates. AS regulates immune cell function independently of transcript abundance (Martinez and Lynch, 2013). In related autoimmune diseases including systemic lupus erythematosus and multiple sclerosis, AS analysis has repeatedly identified disease-relevant gene sets that only partially overlap with those detected by differential expression, exposing candidate mechanisms invisible to expression-only approaches (Papanikolaou et al., 2021; Sak et al., 2025). Direct transcriptome-wide differential AS analysis has not been performed in FMS. Cell-type composition is a well-recognised confounder in bulk tissue transcriptomic analyses (Jaffe and Irizarry, 2014).

Using the female subcohort of GSE221921 (n = 132), we tested for differential AS after adjustment for inferred PBMC composition, alongside parallel differential expression analysis to distinguish splicing-specific from expression-overlapping signals and to identify candidates for future mechanistic follow-up.

## 2. Methods

### 2.1 Data source and study cohort

We analysed the publicly available RNA-sequencing dataset GSE221921 (Mohapatra et al., 2024), comprising 96 FMS patients and 93 healthy controls with peripheral blood mononuclear cell (PBMC) samples. FMS diagnosis in the original study required 2016 American College of Rheumatology criteria (Wolfe et al., 2016) and a positive FM/a cytokine assay. The cohort was strongly imbalanced with respect to sex, and AS varies systematically by sex in blood transcriptomes (Kosmara et al., 2023); we therefore restricted the analysis to female participants (91 FMS, 41 controls; n = 132).

### 2.2 RNA-seq preprocessing and quantification

Raw FASTQ files were retrieved from the Sequence Read Archive (SRP413437). Reads were adapter-removed, trimmed (bases below Phred 20, poly-G tails) and filtered if below 50 nt length using FastQC 0.12.1 and fastp 0.23.4. Reads were aligned to the GRCh38.p14 human reference genome using STAR 2.7.11a, with genes annotated using GENCODE release 44. Reads per gene were normalised to transcripts per million (TPM) for cell-type composition deconvolution (Section 2.3), and gene-level counts derived from the same alignment were used for differential expression analysis (Section 2.5).

### 2.3 Cell-type composition deconvolution

Cell composition of peripheral blood was estimated using the ABIS deconvolution method (Monaco et al., 2019) as implemented in the immunedeconv R package (version 2.1.4, R 4.3.3). ABIS was selected for its PBMC-specific training and high subtype resolution (17 immune cell populations), matching the sample type of GSE221921. TPM values served as input. Composition differences between FMS and controls were tested per cell type with Mann-Whitney U tests; effect sizes are reported as Cliff’s δ. Immature plasma B cells were excluded from the formal multiple-testing analysis (near-zero estimates in both groups); their unadjusted statistics are reported descriptively in Supplementary Table S3. The remaining 16 populations entered Benjamini-Hochberg correction (Benjamini and Hochberg, 1995) at FDR < 0.05. Principal component analysis on the ABIS output yielded composition covariates; the first three principal components were included in the downstream cell type-adjusted differential AS and gene expression analyses.

### 2.4 Differential alternative splicing analysis

AS was analysed with LeafCutter 0.2.9 (Li et al., 2018) using default clustering parameters. Differential AS between FMS and controls was tested with LeafCutter’s covariate-adjusted regression framework, incorporating the first three composition principal components.

Clusters were considered significant at FDR < 0.05 (Benjamini-Hochberg) and max |ΔPSI| ≥ 0.1 across cluster junctions. Significant clusters were subjected to post-hoc quality control by manual inspection in the UCSC Genome Browser and IGV; clusters mapping to genomic regions incompatible with genuine splicing events (unusual intron span in repetitive or ambiguous contexts, or ambiguous assignment to pseudogene-containing regions) were excluded from downstream biological interpretation and are described in Supplementary Materials.

### 2.5 Differential gene expression analysis

Gene-level differential expression was analysed with DESeq2 1.42.0 (Love et al., 2014), incorporating the same three principal components as covariates in addition to disease status. A parallel unadjusted analysis using disease status alone was performed to quantify the composition contribution to the differential expression signal. Genes with an adjusted p-value < 0.05 (Benjamini-Hochberg) and a minimum expression of baseMean ≥ 10 were considered significant.

### 2.6 Ethical considerations

This work is a secondary analysis of publicly available, fully anonymised RNA-sequencing data deposited in the Gene Expression Omnibus under accession GSE221921. The original data collection was approved by the appropriate institutional ethics review body and informed consent was obtained from all participants, as reported in the source publication (Mohapatra et al., 2024). No additional ethics approval was required for this in-silico reanalysis.

## 3. Results

### 3.1 Cohort characteristics

The final analytical female cohort comprised 91 FMS patients and 41 controls (n = 132).

### 3.2 Cell-type composition differs between FMS and controls

Peripheral immune cell composition was estimated across 17 subtypes. One population (immature plasma B cells) was excluded from the formal multiple-testing analysis based on the quality criterion described in Methods; its unadjusted statistics are reported descriptively in Supplementary Table S3. Of the 16 remaining populations, two differed significantly between groups after multiple testing correction (Figure 1A; full statistics in Supplementary Table S3). Conventional monocytes showed the largest effect, with lower ABIS-inferred estimates in FMS (Cliff’s δ = −0.38, FDR = 0.008; Figure 1B). Memory B-cell estimates were higher in FMS (Cliff’s δ = +0.32, FDR = 0.030).

**Figure 1.**
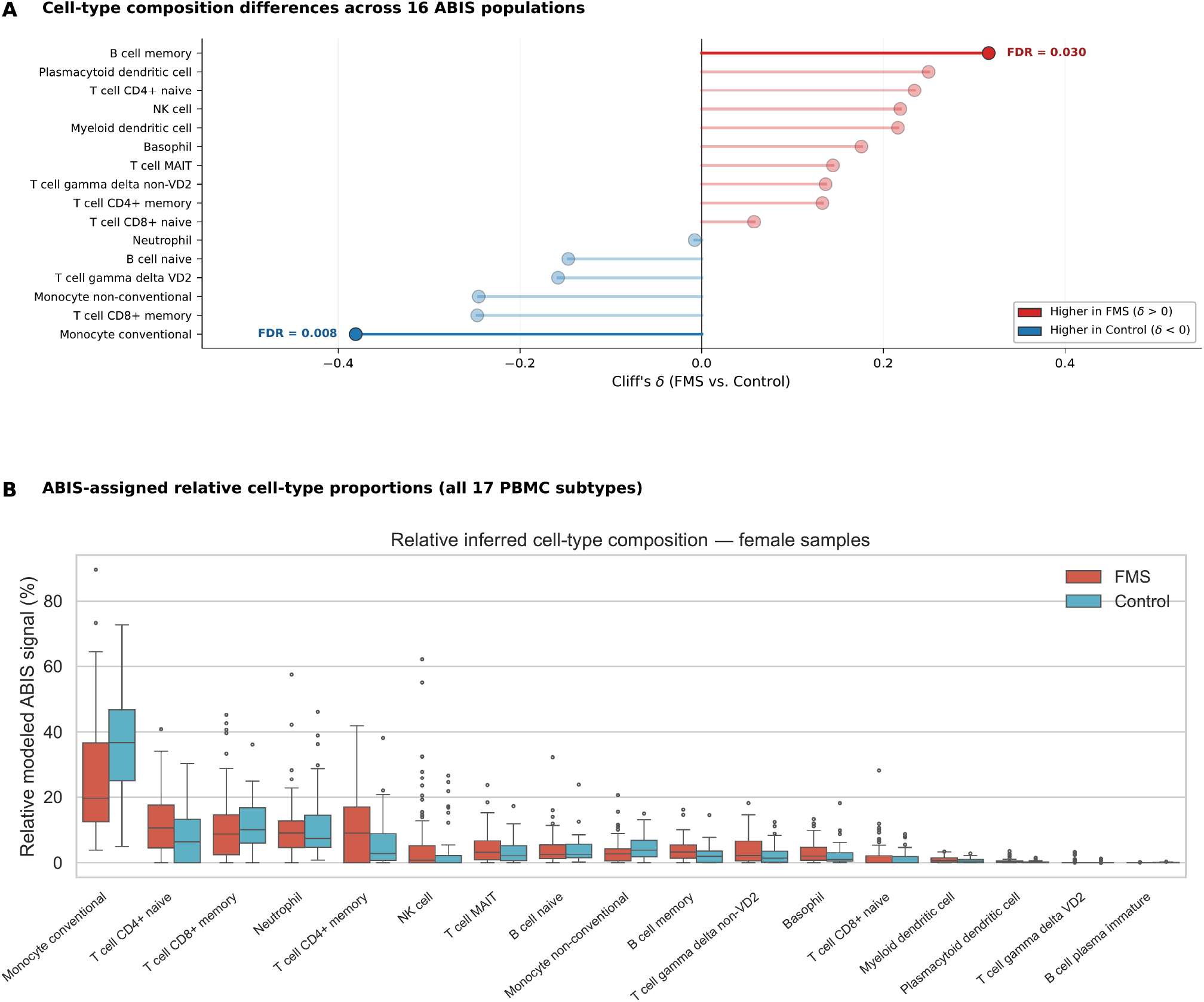
Peripheral immune cell composition in fibromyalgia. (A) Cell-type composition differences between FMS patients and controls across the 16 tested ABIS populations (Cliff’s δ; immature plasma B cells excluded from the formal multiple-testing analysis). Point colour indicates effect direction (red: higher in FMS; blue: higher in controls); populations with FDR < 0.05 are highlighted and labelled. (B) ABIS- inferred cell-type estimates across all 17 PBMC subtypes for FMS (red) and control (blue) samples, scaled to sum to 100 percent for visualisation only; significance tests reported in the main text and Supplementary Table S3 are based on the unscaled signals.

### 3.3 Differential alternative splicing in fibromyalgia

Composition-adjusted LeafCutter analysis identified 14 significant intron clusters. Five were excluded from biological interpretation (three with unusually long or complex junction structures in repetitive or ambiguous genomic context; one with ambiguous locus assignment to a pseudogene-containing region; one — FBXO10 — was not carried forward because of very low gene-level abundance (baseMean = 1.2), limiting robust transcript-level interpretation in this dataset; details in Supplementary Materials). The remaining nine interpretable clusters comprised eight mapping to protein-coding genes (ACAP3, ANKRD28, CR1, GNLY, NVL, REST, RPSA2, TSNARE1) and one to a non-coding locus (ENSG00000242588) (Figure 2A). Effect sizes were substantial, with a median max |ΔPSI| of 0.61 and a maximum of 0.81.

**Figure 2.**
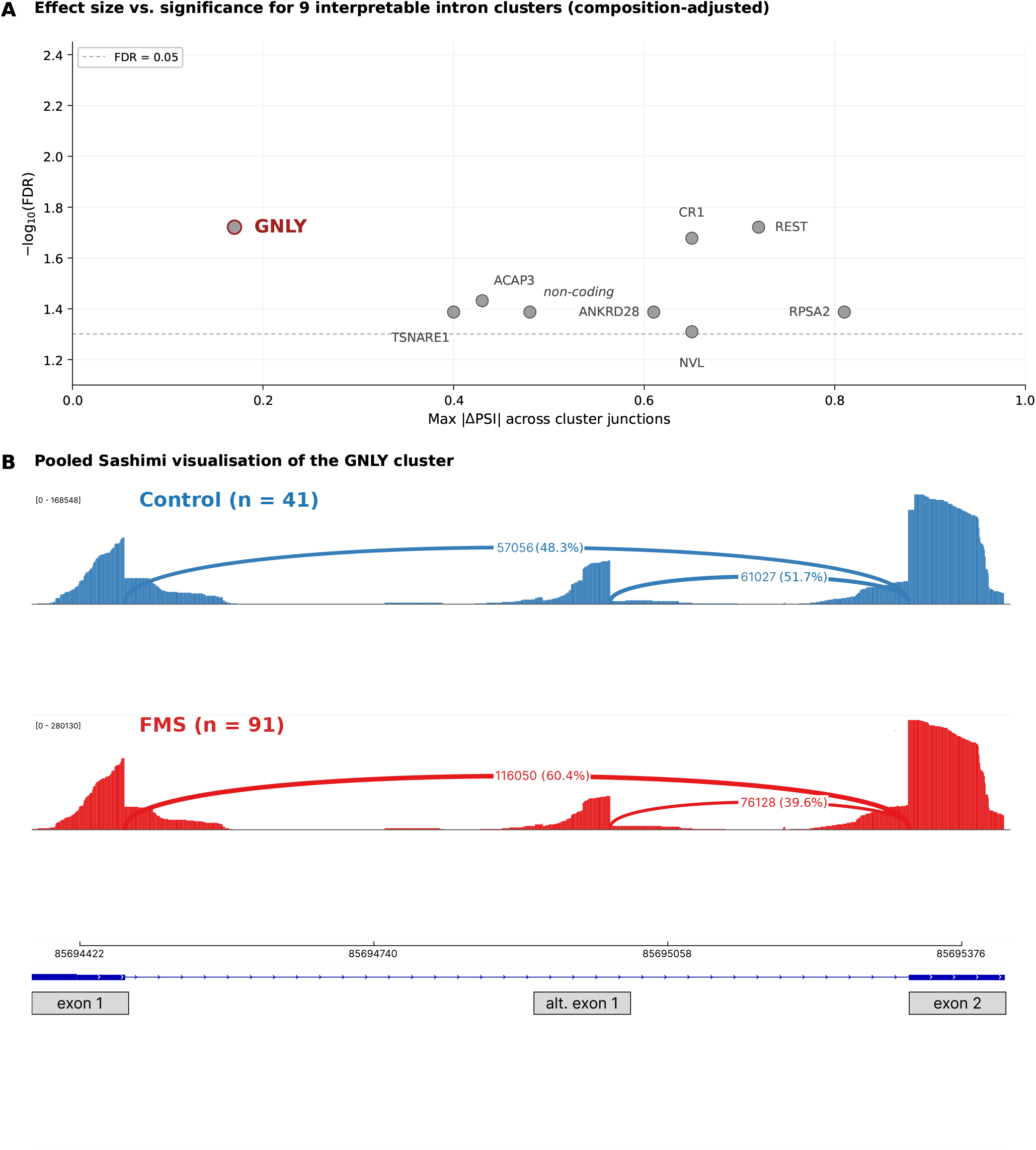
Alternative splicing landscape and GNLY splicing shift in fibromyalgia. (A) Effect size (max |ΔPSI| across cluster junctions) versus significance for the nine interpretable intron clusters (see Table 1; five additional significant clusters were excluded from biological interpretation, four on quality-control grounds and one on grounds of very low gene-level abundance). Eight clusters map to protein-coding genes and one to a non- coding locus. The dashed line marks FDR = 0.05. (B) Pooled Sashimi visualisation of the GNLY cluster (chr2:clu_13300_+). Read coverage and the two competing 5’ splice junctions are shown for Control (n = 41, top, blue) and FMS (n = 91, bottom, red) samples. Numbers on the arcs indicate pooled raw junction read counts; percentages in parentheses indicate the relative usage of the two competing junctions within each group. Coverage tracks are independently scaled and should therefore not be compared quantitatively between groups. The exon 1-exon 2 junction accounted for 48.3 percent of the two junctions in controls and 60.4 percent in FMS, whereas the alt. exon 1-exon 2 junction accounted for 51.7 percent and 39.6 percent, respectively. The composition-adjusted LeafCutter analysis yielded |ΔPSI| = 0.17 (FDR = 0.019).

**Table 1.** Interpretable intron clusters (n = 9) from composition-adjusted LeafCutter analysis.

| Gene | Cluster ID | Junctions | Max $\vert \Delta\text{PSI}\vert $ | Splicing FDR | Expr. $\log_2\text{FC}$ | Expr. padj | Class |
| --- | --- | --- | --- | --- | --- | --- | --- |
| GNLY | chr2:clu_13300_+ | 2 | 0.17 | 0.019 | -0.16 | 0.627 | splicing-specific |
| TSNARE1 | chr8:clu_20673_- | 2 | 0.40 | 0.041 | -0.25 | 0.431 | splicing-specific |
| CR1 | chr1:clu_2211_- | 3 | 0.65 | 0.021 | -0.03 | 0.926 | splicing-specific |
| REST | chr4:clu_16524_+ | 2 | 0.72 | 0.019 | +0.45 | 0.060 | splicing-specific |
| ACAP3 | chr1:clu_1252_- | 2 | 0.43 | 0.037 | -0.59 | 0.019 | shared splicing-expression |
| NVL | chr1:clu_2252_- | 3 | 0.65 | 0.049 | +0.56 | 0.031 | shared splicing-expression |
| ANKRD28 | chr3:clu_15808_- | 2 | 0.61 | 0.041 | +0.58 | 0.032 | shared splicing-expression |
| RPSA2 | chr19:clu_11863_+ | 2 | 0.81 | 0.041 | +0.70 | 0.031 | shared splicing-expression |
| ENSG00000242588 | chr7:clu_20222_+ | 2 | 0.48 | 0.041 | -0.20 | 0.629 | non-coding |

*Junctions: number of alternatively excised introns tested jointly within the cluster. Max $|\Delta\text{PSI}|$ : maximum absolute effect size across cluster junctions. Splicing FDR: cluster-level Benjamini-Hochberg-adjusted p-value from LeafCutter. Expression $\log_2\text{FC}$ and padj from the composition-adjusted DESeq2 analysis. Class: shared splicing-expression, concurrent significant splicing and expression associations (splicing FDR < 0.05 and expression padj < 0.05); splicing-specific, significant splicing without significant differential expression at padj < 0.05; non-coding, cluster mapping to a non-coding locus.*

The GNLY cluster illustrates the junction-level resolution of the analysis (Figure 2B). LeafCutter identified two competing junctions at the 5’ end of the gene, consistent with alternative 5’ splice-junction usage between transcripts with distinct N-terminal sequences. Pooled Sashimi visualisation illustrated the same reciprocal junction-usage pattern between the exon 1-exon 2 junction (chr2:85,694,470–85,695,320) and the alt. exon 1-exon 2 junction (chr2:85,694,996–85,695,320), with the exon 1-exon 2 junction accounting for 60.4 percent of the two competing junctions in FMS versus 48.3 percent in controls (|ΔPSI| = 0.17, FDR = 0.019). GNLY splicing remained associated with FMS after composition adjustment; the corresponding gene-level differential expression was not significant (log2FC = −0.16, padj = 0.627).

### 3.4 Cell composition adjustment reshapes the differential expression landscape

The unadjusted DESeq2 analysis identified 10,803 significant differentially expressed genes. Composition adjustment reduced this number to 3,186 significant genes. Of these, 3,055 remained significant in both analyses, 7,748 lost significance after adjustment, and 131 became significant only after composition adjustment.

### 3.5 Integration of alternative splicing and gene expression distinguishes three FMS-associated categories

Cross-referencing the nine interpretable splicing clusters against the 3,186 composition-adjusted DEGs defines three FMS-associated categories. Four of the eight protein-coding LeafCutter genes are also significantly differentially expressed in the adjusted DEG analysis: ACAP3 (log2FC = −0.59, padj = 0.019), ANKRD28 (+0.58, padj = 0.032), NVL (+0.56, padj = 0.031), and RPSA2 (+0.70, padj = 0.031) (shared splicing–expression). Four further genes (CR1, GNLY, REST, TSNARE1) show significant splicing without significant gene-level differential expression (splicing-specific). The remaining 3,182 significant DEGs represent the expression-specific class, without concurrent significant alternative splicing.

## 4. Discussion

We report the first transcriptome-wide differential AS analysis in FMS and identify GNLY as a splicing-specific immune candidate. This analysis is based on the largest available FMS RNA-sequencing dataset (GSE221921), restricted to the female subcohort (n = 132) and adjusted for inferred PBMC composition.

Among the splicing-specific candidates, GNLY allows the most direct biological interpretation, given the well-characterised function of granulysin as a cytolytic lymphocyte effector with additional extracellular immunomodulatory activity (Krensky and Clayberger, 2009; Tewary et al., 2010). The FMS-associated shift involved alternative 5’ splice-junction usage between transcripts encoding distinct N-terminal sequences, illustrated by pooled Sashimi visualisation (see Results). Because the N terminus can influence protein processing and intracellular targeting, altered GNLY transcript usage may have functional consequences for granulysin handling or secretion, providing a plausible target for functional follow-up in FMS. However, the protein-level consequences of the observed transcript shift cannot be inferred from the present data.

The composition shift we observe is consistent with broader evidence for peripheral immune involvement in FMS. Lower ABIS-inferred conventional monocyte and higher memory B-cell estimates in FMS align with independent reports of altered blood cell ratios in FMS (Kösehasanoğulları et al., 2025). They are also consistent with evidence for B-cell dysregulation in FMS, including passive transfer of pain-related phenotypes by patient IgG (Goebel et al., 2021), anti-satellite glial cell IgG detection (Fanton et al., 2023), and disrupted peripheral B-cell tolerance (Long et al., 2025). Our subtype-resolved estimates identify memory B cells as a specific compartment differing between groups, but we note that ABIS provides inferred cell-type abundance estimates rather than absolute cell counts. The observed differences do not by themselves establish a specific mechanistic link to these immunological phenotypes.

The integrated analysis of AS and gene expression distinguishes three FMS-associated categories. Splicing-specific candidates, exemplified by GNLY and including CR1, REST and TSNARE1, show significant splice-junction associations without significant gene-level differential expression. Shared splicing–expression candidates (ACAP3, ANKRD28, NVL and RPSA2) show concurrent alterations at both molecular levels. Expression-specific signals comprise the majority — 3,182 of the 3,186 composition-adjusted significant DEGs — with no concurrent significant splicing. This distribution parallels findings in systemic lupus erythematosus and multiple sclerosis, where AS captures disease-relevant signal not fully overlapping with differential expression (Papanikolaou et al., 2021; Sak et al., 2025). For FMS specifically, AS identifies candidates not detected by expression analysis alone; the shared class shows concurrent significance at both levels.

Composition adjustment substantially altered the differential expression landscape in this dataset: 7,748 of the 10,803 unadjusted significant genes lost significance after adjustment (approximately 72 percent), and 131 genes became significant only after adjustment. Neither

Mohapatra et al. (2024) nor Zhao et al. (2025) incorporated inferred cell composition as covariates in their transcriptomic models, although the latter estimated cell proportions using CIBERSORT. Mohapatra et al. (2024) reported 1,720 differentially expressed genes in the full mixed-sex cohort; direct numerical comparison with our estimates is confounded by differences in analytical parameters and cohort composition. Beyond gene-level counts, the qualitative message is that a substantial portion of the differential-expression signal in FMS PBMC transcriptomics is composition-sensitive. These findings highlight the need to account explicitly for cell composition when interpreting bulk PBMC transcriptomic studies of FMS, particularly when the aim is to identify composition-independent expression associations.

For clinical FMS research, our findings support two directions. First, peripheral blood transcriptomics remains a productive approach for FMS biomarker discovery, but diagnostic or stratification models built on unadjusted expression signatures risk conflating composition and within-cell regulatory biology, with unclear consequences for cross-cohort generalisability. Second, GNLY provides a candidate for mechanistic follow-up in independent FMS cohorts, ideally with orthogonal validation of the splice-junction shift and assessment of its protein-level or functional consequences. Cross-cohort validation in patients diagnosed by clinical criteria alone (see Limitations) will be important for both directions.

Recent large-scale genetics implicates primarily neuronal biology in FMS (Kerrebijn et al., 2026; Bright et al., 2026); functional evidence supports peripheral immune involvement (Goebel et al., 2021). Our findings add alternative splicing as a regulatory layer within the peripheral immune arm of FMS pathophysiology.

## 5. Limitations

First, the analysis is based on exome-capture RNA-sequencing (SureSelect All Exon V6), which produces uneven read distribution across splice junctions compared to standard total or poly-A RNA-seq protocols. Second, the FMS cohort was selected using the FM/a cytokine assay as an inclusion criterion, defining a specific immunologically characterised subset of FMS rather than the disease as a whole. Findings may not generalise to FMS diagnosed by clinical criteria alone. Third, this is a cross-sectional analysis and does not permit causal inference.

Fourth, our biological interpretation of individual candidate genes rests on a small number of retained clusters, and confirmation in independent cohorts remains essential. Fifth, individual-level age data were not available in the publicly accessible GEO metadata and could therefore not be included as a covariate. Finally, our composition adjustment relies on inferred rather than directly measured cell abundance, so residual composition confounding cannot be entirely ruled out, particularly for cell subsets not well resolved by ABIS deconvolution.

## Acknowledgements

The authors thank the investigators of the original GSE221921 study for making their data publicly available.

## Data availability

RNA-sequencing data are publicly available from the Gene Expression Omnibus under accession GSE221921. Junction-level AS results, gene-level differential expression results, and ABIS deconvolution statistics are provided in Supplementary Tables S1–S3. Analysis code and additional processed data are available from the corresponding author upon reasonable request.

## Author contributions

YB conceived the study, coordinated the analysis strategy, interpreted results, and drafted the manuscript. AN performed the bioinformatic pipeline, developed the covariate-adjusted analysis framework, and contributed to result interpretation and manuscript revision. Both authors have read and approved the final manuscript.

## References

1. Benjamini Y, Hochberg Y. Controlling the false discovery rate: a practical and powerful approach to multiple testing. J R Stat Soc Ser B. 1995;57(1):289–300.

2. Bright U, Beck S, Levey DF, et al. The genetics of fibromyalgia and its relationships to psychiatric and medical traits. Nat Commun. 2026;17:6248. doi:10.1038/s41467-026-75256-6

3. D’Souza RS, Klasova J, Morsi M, et al. The prevalence of fibromyalgia in the general population and at-risk subpopulations: a systematic review and meta-analysis. Anesth Analg. 2026. doi:10.1213/ANE.0000000000008098

4. Fanton S, Menezes J, Krock E, et al. Anti-satellite glia cell IgG antibodies in fibromyalgia patients are related to symptom severity and to metabolite concentrations in thalamus and rostral anterior cingulate cortex. Brain Behav Immun. 2023;114:371–382. doi:10.1016/j.bbi.2023.09.003

5. Goebel A, Krock E, Gentry C, et al. Passive transfer of fibromyalgia symptoms from patients to mice. J Clin Invest. 2021;131(13):e144201. doi:10.1172/JCI144201

6. Häuser W, Ablin J, Fitzcharles MA, et al. Fibromyalgia. Nat Rev Dis Primers. 2015;1:15022. doi:10.1038/nrdp.2015.22

7. Häuser W, Perrot S, Clauw DJ, Fitzcharles MA. Unravelling fibromyalgia: steps toward individualized management. J Pain. 2018;19(2):125–134. doi:10.1016/j.jpain.2017.08.009

8. Heidari F, Afshari M, Moosazadeh M. Prevalence of fibromyalgia in general population and patients, a systematic review and meta-analysis. Rheumatol Int. 2017;37(9):1527–1539. doi:10.1007/s00296-017-3725-2

9. Jaffe AE, Irizarry RA. Accounting for cellular heterogeneity is critical in epigenome-wide association studies. Genome Biol. 2014;15(2):R31. doi:10.1186/gb-2014-15-2-r31

10. Kerrebijn I, Bjornsdottir G, Arbabi K, et al. The genetic architecture of fibromyalgia across 2.5 million individuals. Nat Med. 2026 Jul 28. doi:10.1038/s41591-026-04492-6

11. Kösehasanoğulları M, Aygün Bilecik N, Büyükvural Şen S, Koçyiğit BF. Elevated monocyte-to-lymphocyte and platelet-to-lymphocyte ratios are associated with disease activity and pain in fibromyalgia: a cross-sectional study. J Clin Med. 2025;15(1):155. doi:10.3390/jcm15010155

12. Kosmara D, Papanikolaou S, Nikolaou C, Bertsias G. Extensive alternative splicing patterns in systemic lupus erythematosus highlight sexual differences. Cells. 2023;12(23):2678. doi:10.3390/cells12232678

13. Krensky AM, Clayberger C. Biology and clinical relevance of granulysin. Tissue Antigens. 2009;73(3):193–198. doi:10.1111/j.1399-0039.2008.01218.x

14. Li YI, Knowles DA, Humphrey J, et al. Annotation-free quantification of RNA splicing using LeafCutter. Nat Genet. 2018;50(1):151–158. doi:10.1038/s41588-017-0004-9

15. Long A, Choi Chiu A, Onupom O, et al. Defective peripheral B cell tolerance leads to dysregulated B cell responses in Fibromyalgia Syndrome. bioRxiv. 2025. doi:10.64898/2025.12.16.694591

16. Love MI, Huber W, Anders S. Moderated estimation of fold change and dispersion for RNA-seq data with DESeq2. Genome Biol. 2014;15(12):550. doi:10.1186/s13059-014-0550-8

17. Martinez NM, Lynch KW. Control of alternative splicing in immune responses: many regulators, many predictions, much still to learn. Immunol Rev. 2013;253(1):216–236. doi:10.1111/imr.12047

18. Mohapatra G, Dachet F, Coleman LJ, Gillis B, Behm FG. Identification of unique genomic signatures in patients with fibromyalgia and chronic pain. Sci Rep. 2024;14(1):3949. doi:10.1038/s41598-024-53874-8

19. Monaco G, Lee B, Xu W, et al. RNA-Seq signatures normalized by mRNA abundance allow absolute deconvolution of human immune cell types. Cell Rep. 2019;26(6):1627–1640.e7. doi:10.1016/j.celrep.2019.01.041

20. Papanikolaou S, Bertsias GK, Nikolaou C. Extensive changes in transcription dynamics reflected on alternative splicing events in systemic lupus erythematosus patients. Genes (Basel). 2021;12(8):1260. doi:10.3390/genes12081260

21. Sak M, Chariker JH, Rouchka EC. Systematic analysis of alternative splicing in transcriptomes of multiple sclerosis patient brain samples. Int J Mol Sci. 2025;26(17):8195. doi:10.3390/ijms26178195

22. Tewary P, Yang D, de la Rosa G, et al. Granulysin activates antigen-presenting cells through TLR4 and acts as an immune alarmin. Blood. 2010;116(18):3465–3474. doi:10.1182/blood-2010-03-273953

23. Wolfe F, Clauw DJ, Fitzcharles MA, et al. 2016 Revisions to the 2010/2011 fibromyalgia diagnostic criteria. Semin Arthritis Rheum. 2016;46(3):319–329. doi:10.1016/j.semarthrit.2016.08.012

24. Zhao F, Zhao J, Li Y, et al. Identification of diagnostic biomarkers for fibromyalgia using gene expression analysis and machine learning. Front Genet. 2025;16:1535541. doi:10.3389/fgene.2025.1535541

