## Supplementary Materials for "Transcriptome-wide analysis of alternative splicing in women with fibromyalgia highlights GNLY as an immune candidate"

**Contents:**

Supplementary Note S1. Post-hoc quality control of LeafCutter clusters

Supplementary Table S1. Junction-level statistics for the nine interpretable intron clusters (plus FBXO10, retained for completeness)

Supplementary Table S2. Complete gene-level differential expression results (separate .xlsx file)

Supplementary Table S3. ABIS-inferred cell composition statistics for all 17 immune populations

**Supplementary Note S1. Post-hoc quality control of LeafCutter clusters**

The composition-adjusted LeafCutter analysis identified 14 significant intron clusters (FDR < 0.05, max |ΔPSI| ≥ 0.1). Post-hoc manual review in the UCSC Genome Browser and IGV (see Methods, Section 2.4) retained 9 clusters for biological interpretation and excluded 5 from downstream interpretation, as follows.

Three clusters (chr3:clu_15335_+, chr21:clu_14250_+, chr21:clu_14416_-) mapped to unusually long or complex junction structures in repetitive or ambiguous genomic context, inconsistent with typical alternative splicing events. A fourth cluster (chr5:clu_17257_-) mapped to a region overlapping the annotated pseudogene GUSBP3 and additional Ensembl gene identifiers, precluding unambiguous assignment to a single biologically interpretable gene locus.

A fifth cluster (chr9:clu_21621_-) mapped to FBXO10, a gene with very low overall expression in this dataset (baseMean = 1.2, well below the abundance threshold used for the differential expression analysis). The junction-level statistics for this cluster are retained in Supplementary Table S1 for completeness but were not carried forward for biological interpretation, given that transcript-level inferences from a locus with this level of abundance are difficult to interpret robustly.

**Supplementary Table S1. Junction-level statistics for the nine interpretable intron clusters (plus FBXO10, retained for completeness) from composition-adjusted LeafCutter analysis (n = 132).**

| **Gene** | **Cluster ID** | **Junction coordinates (GRCh38)** | **Strand** | **ΔPSI** | **p-value** | **FDR** | **Classification** |
| --- | --- | --- | --- | --- | --- | --- | --- |
| **GNLY** | chr2:clu_13300_+ | chr2:85,694,470–85,695,320 | + | −0.168 | 1.0×10⁻⁵ | 0.019 | Splicing-specific |
| GNLY | chr2:clu_13300_+ | chr2:85,694,996–85,695,320 | + | +0.168 | 1.0×10⁻⁵ | 0.019 | Splicing-specific |
| TSNARE1 | chr8:clu_20673_- | chr8:142,271,691–142,273,057 | − | +0.403 | 4.5×10⁻⁵ | 0.041 | Splicing-specific |
| TSNARE1 | chr8:clu_20673_- | chr8:142,271,691–142,274,781 | − | −0.403 | 4.5×10⁻⁵ | 0.041 | Splicing-specific |
| CR1 | chr1:clu_2211_- | chr1:207,552,851–207,558,538 | − | +0.655 | 1.1×10⁻⁵ | 0.021 | Splicing-specific |
| CR1 | chr1:clu_2211_- | chr1:207,552,851–207,575,595 | − | −0.322 | 1.1×10⁻⁵ | 0.021 | Splicing-specific |
| CR1 | chr1:clu_2211_- | chr1:207,569,946–207,575,595 | − | −0.333 | 1.1×10⁻⁵ | 0.021 | Splicing-specific |
| REST | chr4:clu_16524_+ | chr4:56,908,213–56,910,630 | + | −0.716 | 1.3×10⁻⁵ | 0.019 | Splicing-specific |
| REST | chr4:clu_16524_+ | chr4:56,909,039–56,910,630 | + | +0.716 | 1.3×10⁻⁵ | 0.019 | Splicing-specific |
| ACAP3 | chr1:clu_1252_- | chr1:1,298,113–1,298,303 | − | +0.431 | 3.2×10⁻⁵ | 0.037 | Shared splicing–expression |
| ACAP3 | chr1:clu_1252_- | chr1:1,298,113–1,298,370 | − | −0.431 | 3.2×10⁻⁵ | 0.037 | Shared splicing–expression |
| NVL | chr1:clu_2252_- | chr1:224,300,663–224,301,691 | − | −0.409 | 8.1×10⁻⁵ | 0.049 | Shared splicing–expression |
| NVL | chr1:clu_2252_- | chr1:224,300,663–224,303,723 | − | +0.647 | 8.1×10⁻⁵ | 0.049 | Shared splicing–expression |
| NVL | chr1:clu_2252_- | chr1:224,301,771–224,303,723 | − | −0.238 | 8.1×10⁻⁵ | 0.049 | Shared splicing–expression |
| ANKRD28 | chr3:clu_15808_- | chr3:15,795,306–15,797,971 | − | −0.580 | 5.3×10⁻⁵ | 0.041 | Shared splicing–expression |
| ANKRD28 | chr3:clu_15808_- | chr3:15,795,306–15,859,377 | − | +0.613 | 5.3×10⁻⁵ | 0.041 | Shared splicing–expression |
| RPSA2 | chr19:clu_11863_+ | chr19:23,807,939–23,808,045 | + | −0.808 | 4.6×10⁻⁵ | 0.041 | Shared splicing–expression |
| RPSA2 | chr19:clu_11863_+ | chr19:23,808,154–23,808,719 | + | +0.808 | 4.6×10⁻⁵ | 0.041 | Shared splicing–expression |
| *ENSG00000242588* | chr7:clu_20222_+ | chr7:128,642,281–128,647,923 | + | −0.482 | 5.4×10⁻⁵ | 0.041 | Non-coding |
| *ENSG00000242588* | chr7:clu_20222_+ | chr7:128,642,281–128,648,792 | + | +0.482 | 5.4×10⁻⁵ | 0.041 | Non-coding |
| *FBXO10* | *chr9:clu_21621_-* | *chr9:37,541,774–37,576,211* | *−* | *+0.165* | *7.3×10⁻⁵* | *0.048* | *Not carried forward** |
| *FBXO10* | *chr9:clu_21621_-* | *chr9:37,541,774–37,588,811* | *−* | *−0.165* | *7.3×10⁻⁵* | *0.048* | *Not carried forward** |

*Junction-level detail for the intron clusters. Each cluster contributes two or three competing junctions; the signs of the ΔPSI values indicate junction-level (not cluster-level) direction of change. Junction coordinates are 1-based (GRCh38). p-value and FDR are cluster-level statistics from LeafCutter and are therefore identical for all junctions within a cluster. Classification: splicing-specific, significant splicing without significant gene-level differential expression at padj < 0.05; shared splicing–expression, concurrent significant splicing and expression associations (splicing FDR < 0.05 and expression padj < 0.05); non-coding, cluster mapping to a non-coding locus; not carried forward*, cluster retained here for completeness of the LeafCutter output but not carried forward for biological interpretation because of very low gene-level abundance (baseMean = 1.2), limiting robust transcript-level interpretation.*

**Supplementary Table S3. ABIS-inferred cell composition statistics for all 17 immune populations (n = 132).**

| **Immune cell population** | **FMS median (IQR)** | **Control median (IQR)** | **Cliff's δ** | **p-value** | **FDR** |
| --- | --- | --- | --- | --- | --- |
| **Monocyte conventional** | 3.170 (1.700–4.900) | 5.970 (3.370–8.830) | −0.381 | 4.8×10⁻⁴ | 0.008 |
| **B cell memory** | 0.594 (0.149–1.025) | 0.297 (0.040–0.554) | +0.316 | 3.8×10⁻³ | 0.030 |
| Plasmacytoid dendritic cell | 0.038 (0.013–0.108) | 0.024 (0.009–0.056) | +0.250 | 2.2×10⁻² | 0.078 |
| T cell CD8+ memory | 1.120 (0.273–1.940) | 1.540 (0.992–2.920) | −0.247 | 2.4×10⁻² | 0.078 |
| Monocyte non-conventional | 0.355 (0.048–0.750) | 0.703 (0.266–1.020) | −0.246 | 2.5×10⁻² | 0.078 |
| T cell CD4+ naive | 1.750 (0.362–3.360) | 0.661 (−0.006–2.200) | +0.234 | 3.2×10⁻² | 0.085 |
| NK cell | 0.097 (0.001–0.877) | 0.008 (−0.059–0.270) | +0.219 | 4.5×10⁻² | 0.096 |
| Myeloid dendritic cell | 0.122 (0.026–0.307) | 0.086 (0.012–0.149) | +0.216 | 4.8×10⁻² | 0.096 |
| Basophil | 0.331 (0.086–1.044) | 0.179 (0.073–0.587) | +0.176 | 1.1×10⁻¹ | 0.191 |
| T cell gamma delta VD2 | −0.112 (−0.290–−0.018) | −0.069 (−0.199–−0.017) | −0.158 | 1.5×10⁻¹ | 0.236 |
| B cell naive | 0.389 (0.124–0.788) | 0.501 (0.246–0.964) | −0.147 | 1.8×10⁻¹ | 0.248 |
| T cell MAIT | 0.559 (0.110–1.305) | 0.409 (0.113–0.907) | +0.144 | 1.9×10⁻¹ | 0.248 |
| T cell gamma delta non-VD2 | 0.295 (0.064–1.330) | 0.300 (0.040–0.708) | +0.136 | 2.1×10⁻¹ | 0.255 |
| T cell CD4+ memory | 1.080 (−0.030–3.125) | 0.456 (0.073–1.940) | +0.133 | 2.2×10⁻¹ | 0.255 |
| T cell CD8+ naive | −0.384 (−0.931–0.452) | −0.377 (−1.080–0.252) | +0.058 | 6.0×10⁻¹ | 0.639 |
| Neutrophil | 1.380 (0.576–2.370) | 1.160 (0.568–2.500) | −0.008 | 9.4×10⁻¹ | 0.945 |
| *B cell plasma immature* | *0.000 (−0.002–0.004)* | *0.005 (−0.001–0.019)* | *−0.350* | *1.3×10⁻³* | *n.t.†* |

*Full ABIS deconvolution statistics for the female subcohort of GSE221921 (91 FMS, 41 controls). Values are ABIS-inferred cell-population estimates (arbitrary units from the immunedeconv 2.1.4 output; not scaled to sum to 100 percent). Effect sizes are Cliff's δ (negative: lower in FMS; positive: higher in FMS); p-values from two-sided Mann-Whitney U tests. FDR was computed by Benjamini-Hochberg across the 16 tested populations. Populations are sorted by absolute effect size; the two populations reaching FDR < 0.05 are highlighted in bold. † Immature plasma B cells were excluded from the formal multiple-testing analysis (near-zero estimates in both groups); their unadjusted statistics are shown descriptively. Figure 1B displays the same underlying data rescaled to sum to 100 percent for visualisation only; significance tests reported in the main text and in this table are based on the unscaled signals.*
